# Fluctuations and entropy enable neural crest cell ingression

**DOI:** 10.1101/2023.02.10.528070

**Authors:** Clarissa C. Pasiliao, Evan C. Thomas, Theodora Yung, Min Zhu, Hirotaka Tao, Yu Sun, Sidhartha Goyal, Sevan Hopyan

**Affiliations:** Program in Developmental and Stem Cell Biology, Research Institute, Hospital for Sick Children, Toronto, ON, Canada M5G 0A4; Department of Molecular Genetics, University of Toronto, M5S 1A8; Department of Mechanical and Industrial Engineering, University of Toronto, M5S 3G8; Department of Physics, University of Toronto, M5S 1A7; Division of Orthopaedic Surgery, Hospital for Sick Children and University of Toronto, M5G 1X8

**Author notes:** Correspondence: Sevan Hopyan, 686 Bay Street, 16-9713, Toronto, Ontario M5G 0A4, Canada. Entropic cell ingression.

## Abstract

The second law of thermodynamics explains the dissipative nature of embryonic development as an exchange of energy-dependent order for proportionately greater output of heat and waste. Recent work on granular matter provides a path by which to define the roles of passive, stochastic mechanisms in nonequilibrium systems. Here, we apply such a framework to examine the role of thermodynamic parameters to cell ingression, the movement of cells from one tissue layer to another that has been attributed, in part, to directional cues. Using the murine neural crest as a model system, we provide evidence that a stochastic mechanism, rather than a proposed stiffness gradient, underlies cell ingression. Cortical fluctuations representing effective temperature and cell packing configurations generate an entropic trap that promotes cell ingression. The results imply dissipative mechanisms that transiently disorder tissue underlie some morphogenetic events.

## INTRODUCTION

At first glance, embryonic development seems to defy the second law of thermodynamics due to its progressively organising nature. However, living entities are open systems that use and transform energy into metabolic waste and heat to an extent that more than compensates for the order they temporarily exhibit. As such, embryos are dissipative because they function far from thermodynamic equilibrium^1-4^. Interestingly, progression towards greater overall structural order during development is punctuated by periods of disorder^5^, suggesting that stochastic processes contribute to morphogenesis.

Cell ingression is a fundamental developmental process in which cells move from one tissue layer to another as in epithelial to mesenchymal transition^6-8^. This process occurs at multiple sites in the embryo to deliver specialised cells to organ primordia^9-12^. A well-known example is the ingression of cells from the neural crest into cranial mesoderm from where they subsequently migrate to craniofacial regions and differentiate into various cell types^13-15^. The specification^16^, differentiation potential^17,18^ and migration^14,19^ of neural crest cells are increasingly understood. Proposals to explain the ingression event often invoke intriguing biochemical and physical cues^20-33^ but may be incomplete since stochastic possibilities are usually not considered. Here, we apply a framework that enables thermodynamic assessment of nonequilibrium systems using effective temperature^34^ in combination with empirical biophysical measurements in the mouse embryo. We find cortical fluctuations provide stochastic kicks that are essential for crossing an energy-like barrier and cell packing configurations generate an entropic trap that enables spontaneous neural crest cell ingression.

## RESULTS

### Cell ingression in the murine neural crest

In various systems, ingressing cells exhibit stereotypical bottle shape morphology through apical constriction that is mediated by contractions of non-muscle myosin II, cell rounding and blebbing^35-42^. However, these processes are not consistent between zebrafish and chick embryos^43,44^ and have not been detailed in the mouse embryo. To visualise neural crest cell ingression, we used time-lapse lightsheet imaging of live, intact Wnt1:Cre2^Tg/+^;mT/mG mouse embryos at embryonic day E8.5. For orientation, the 12-somite embryo demonstrates the neural crest and ingressed cells transiting to facial primordia (Mov. S1). We first observed evidence of cell delamination within the midbrain of 4-somite embryos among rosette-like cell clusters that were separated by 1-10 cell diameters (Fig. 1A, Mov. S2). The central, leading cell of a cluster descended basally as neighbouring cells moved in to fill the gap, akin to T2 transitions in vertex models of epithelia^45^ (Fig. 1B, C, Mov. S3, S4). Ingressing cell clusters entered mesoderm in separate, stalactite-like chains (Fig. 1D, Mov. S5) that gradually coalesced (Fig. S1A, Mov. S6) into a coherent subepithelial collection (Fig. S1B). Subsequently, ingressed cells converged and extended into a cone-like configuration as they migrated deeper (Mov. S7). We focused further on mechanisms underlying the initial ingression.

**Figure 1.**
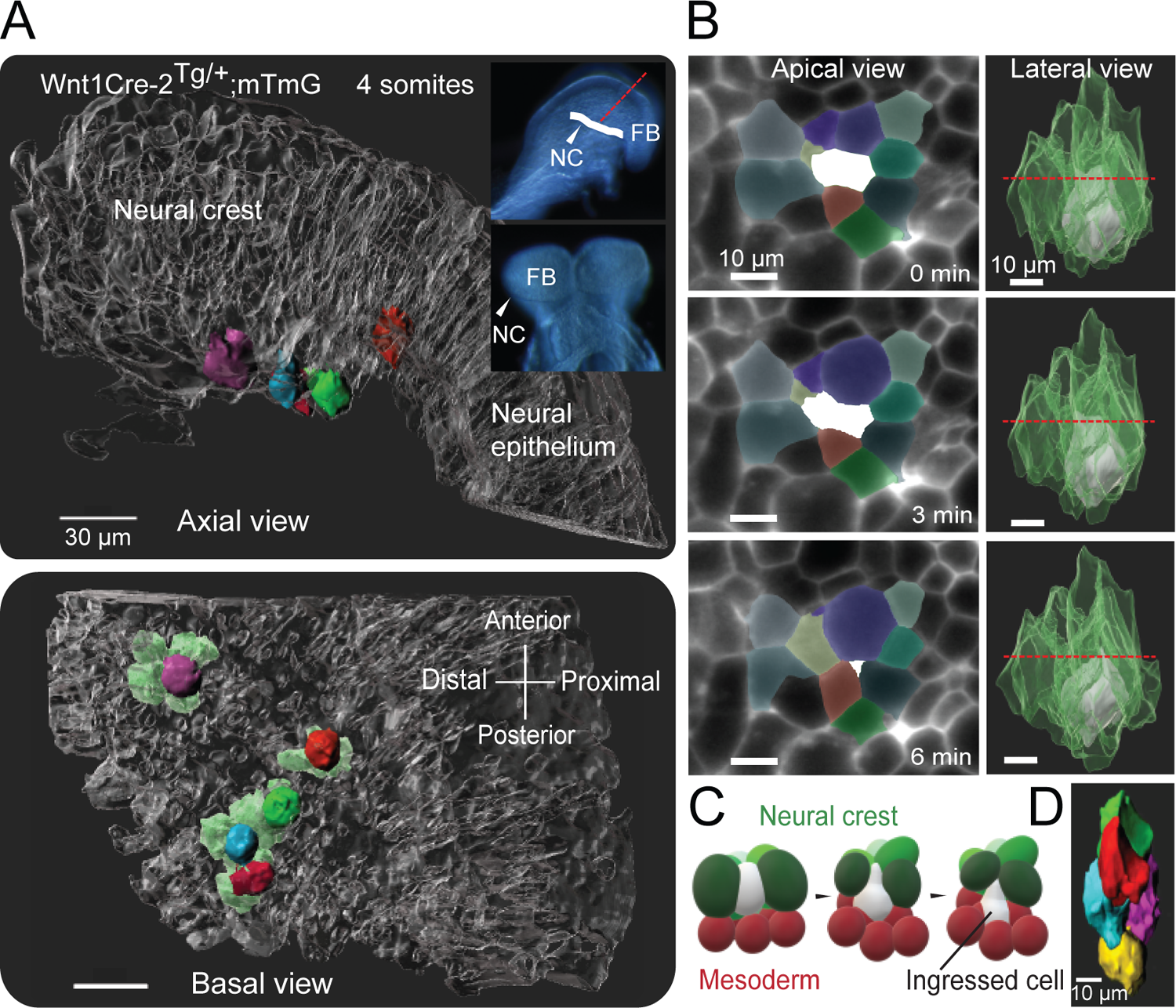
Neural crest cells ingress through a series of T2 transitions. **A** Cell ingression at the cranial neural crest of a 4-somite *Wnt1:Cre-2*^*Tg/+*^*;mT/mG* embryo. Individual lead cells are marked by solid colours while neighbouring, pre-ingressed cells are highlighted in light green in the lower panel. Inset: Intact mouse embryo, 4 somite stage. Dashed red line indicates the axial level shown in the top panel. White line indicates the neural crest (NC). Bottom panel: basal/mesodermal, or deep to superficial, view of ingressing leader cells. Images are representative of 3 embryos at 4 somite stage. FB indicates forebrain. Scale bars represent 30 µm. **B** Apical and whole cell rendering of a T2 transition within the neural crest. Left panels: epithelial planar sections showing the apical area of the central cell (white) and immediate neighbours in the same plane. Right panels: three-dimensional renderings of the central white cell and its neighbours (green). Images are representative of 3 embryos at 5-6 somite stage. Scale bars represent 10 µm. **C** Schematic of neural crest ingression. As a neural crest cell ingresses (white) into the mesoderm (red), its apical surface diminishes and the neighbouring cells (green) move in to occupy the vacated space. **D** Rendering of actual neural crest cells within an ingressing ‘stalactite’. Lead cell (yellow) is closely followed by a chain of neural crest cells moving into the mesoderm.

### A thermodynamic perspective of cell ingression

Advances toward a biophysical explanation for neural crest cell ingression that complement descriptions of cellular morphology changes include evidence for permissive and inductive mechanisms. Proposals include transition of the repertoire of cadherins that regulate cell adhesion affinities from epithelial to mesodermal^20^. However, the extent to which cadherin changes are required for ingression versus subsequent migration is unclear, and at least some transcriptional regulators of cadherins are dispensable for neural crest cell ingression in the mouse^46,47^. Planar cell polarity signalling is required for neural crest cell ingression in *Xenopus* and zebrafish^21,22^, but VANGL-dependent PCP is not required in the mouse^48,49^. Discontinuities of the basement membrane^23-32^ may be necessary but are not sufficient for neural crest cell ingression^27,28^. An inductive cue proposed in *Xenopus* is an increase in mesodermal stiffness that triggers ingression of neural crest cells downstream of PCP signalling^33^. In the zebrafish myocardium, epithelial crowding underlies tensile heterogeneity among cells^50^ and, in conceptually related work in *Xenopus*, it was shown that downregulation of cellular tension promotes ingression^51^. Interestingly, enhanced aerobic glycolysis is required at the onset of neural crest cell ingression in the chick, and the expression of genes encoding rate-limiting glycolytic enzymes diminishes among post-ingression migratory cells^52^. We infer from these advances that mechanical factors drive and constrain ingression and cells likely overcome an energy barrier to delaminate, although a cohesive mechanism remains elusive.

The role of stiffness as an ingression cue is unclear because surface indentation methods in association with epithelial elevation used in *Xenopus* are not optimal for measuring subsurface stiffness gradients^33,53^. To measure stiffness in the mouse embryo, we used a 3D magnetic tweezer system that is suitable for bulk mesoderm without the need for dissection^54^. Stiffness of the neural crest was comparable to that of the underlying mesoderm but increased linearly beginning at a distance of 60-80 µm deep to the basement membrane (Fig. S1C). Therefore, durotaxis cannot precipitate ingression in the mouse but may contribute to deeper migration as has been suggested in *Xenopus*^53^. Although other directional cues may be present, we questioned whether a more stochastic process might suffice for cell ingression.

Conjecture and observations have been applied to describe evolution and embryonic development as dissipative processes^1,2,4,55^. However, it is not clear whether periods of disorder contribute to specific developmental events. Buoyed by our observations of cortical fluctuations among ingressing cells (more on this below), we considered the possibility of fluctuation-driven transitions as a driver of cell ingression. This concept is akin to increased effective temperature facilitating barrier crossing in equilibrium systems (Fig. S2A).

To identify the level of stochastic fluctuations for cells to ingress, we created a two-dimensional vertex model of cell ingression that employed active cell shape fluctuations which are essential for cell intercalation^37,56,57^ and passive, dissipative relaxation that is central to any non-equilibrium setting (Fig. 2A, Methods). To relate this model to work on non-equilibrium granular systems^58,59^, we showed that increased cortical fluctuations in the model lead to an increase in the effective temperature^34^ (Fig. 2B, S2B, S2C, Methods). Compared to a deterministic path in which junctions were rearranged to force cells beyond an energy barrier, fluctuations resulted in flattening of the energy landscape. “Hot” cells moved freely into mesoderm with effectively no barrier (Fig. 2C).

**Figure 2.**
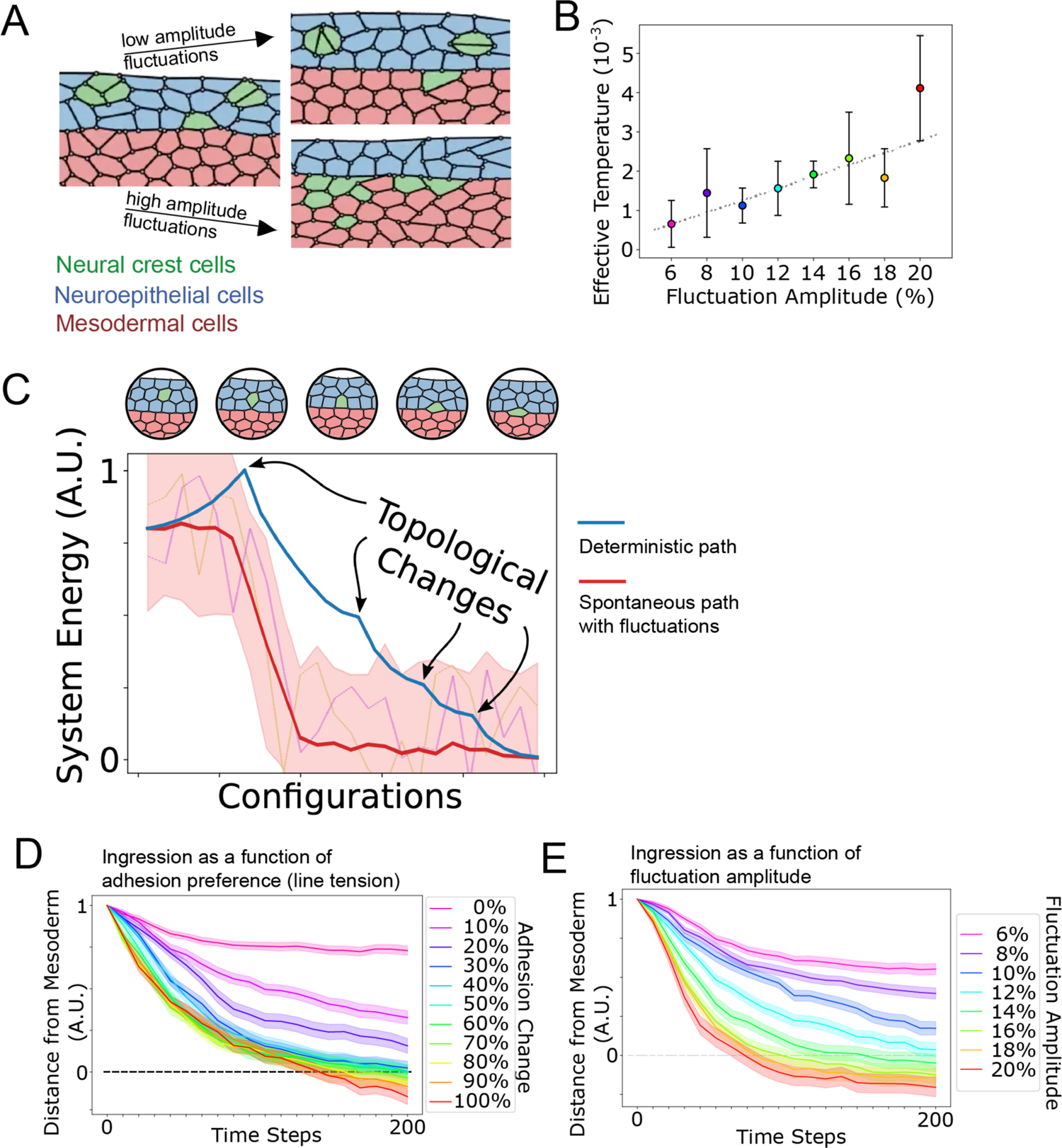
Simulated effect of cell fluctuations. **A** The initial and final states of vertex model simulations in which neural crest cells (green) move spontaneously from the epithelium (blue) toward mesoderm (red) as a function of fluctuation amplitude. **B** Teff, given by the ratio of particle diffusivity (D) to susceptibility (χ), was positively correlated with amplitude of cortical fluctuations in simulations (n=48 simulations per amplitude). **C** Plot of system energy versus configurations of trajectories for single cell ingressions. The blue curve represents a deterministic path wherein the closest edge separating an ingressing cell from the mesenchyme is contracted to a point manually before the system is allowed to relax in the absence of fluctuations. That curve shows there is an energy barrier that must be overcome for cells to ingress into mesoderm. The red curve is the average of 96 simulations starting from the same initial configuration but with stochastic cortical fluctuations added and without manual interference (standard deviation shaded). The curves were shifted to normalise the time step of ingression. Two representative curves are shown as orange and purple dotted curves. **D** Rates of ingression as a function of adhesion affinity. Intercellular line tensions were unchanged (0%) in cells that maintained their epithelial character. To mimic the acquisition of mesodermal adhesion character as cells undergo epithelial-to-mesenchymal transition, line tensions were altered to partially resemble that of mesoderm from (e.g. blue curve represents line tension of 20% mesodermal and 80% epithelial). Cortical fluctuations were maintained at 12% (as in Fig. 2E). **E** Rates of ingression as a function of cortical fluctuation amplitudes. Amplitudes are represented as the standard deviation (% of the mean) of cell shape fluctuations. Adhesion preference for mesoderm was set at 60% (as in Fig. 2D). For D and E, solid colour curves represent the mean (of means of cell positions) and shading represents standard error (sigma/square root(n)) of 48 simulations for each condition.

One of the most extensively documented biophysical parameters in model organisms including the mouse is expression of mesodermal cadherins by neural crest cells prior to ingression, thereby increasing their sorting preferences for mesoderm^18,46,60-68^. As expected, when simulated neural crest cell adhesion affinities were increasingly changed from epithelial to mesodermal (represented by changing intercellular line tensions^69^ in the model), ingression of hot cells into mesoderm became correspondingly more efficient (Fig. 2D).

When simulated neural crest cell fluctuation amplitudes were low, cell movements toward mesoderm were relatively slow and incomplete with some cells becoming trapped within the epithelial layer despite a high adhesion preference for mesoderm (60% as in Fig. 2D). Conversely, when fluctuation amplitudes were high, ingression occurred more rapidly and completely (Fig. 2A, E, Mov. 8, 9). These simulations suggest that adhesion preferences alone are insufficient and that relatively high amplitude cortical fluctuations could drive stochastic cell ingression.

### Biophysical parameters in the murine neural crest

To test our model predictions, we recognised that internal energy (U) cannot be measured *in vivo* because it includes numerous, immeasurable entities such as molecular bond energies. However, it is possible to determine whether the entropic difference between pre-ingressed and ingressed neural crest cells may enable their ingression. We examined changes in configurational entropy as a function of packing which influences the collective behaviours of athermal materials^70^ and living tissues^57,71^ alike. At the pre-ingressed neural crest, cells were more densely packed than in the neighbouring pseudostratified columnar neuroepithelium and in mesoderm (Fig. 3A). Interstitial volumes were substantially greater in mesoderm relative to the neuroepithelium and pre-ingressed neural crest (Fig. 3B). Together, these data indicate that pre-ingressed and ingressed neural crest cells represent the most and least physically constrained populations, respectively. To define cell shape and neighbour relationships, we segmented 3D volumes (Fig. S2D, Methods). The non-parametric cell shape index (S/V^2/3^) was progressively lower, indicating that cells were rounder, between the neuroepithelium, pre-ingressed neural crest and ingressed neural crest cells (Fig. 3C, D). The number of cell neighbours increased from a median of 7 in the neuroepithelium, to 10 among pre-ingressed neural crest cells and 12 among ingressed cells (Fig. 3E). Therefore, ingressing neural crest cells transition to a progressively unconstrained state,

**Figure 3.**
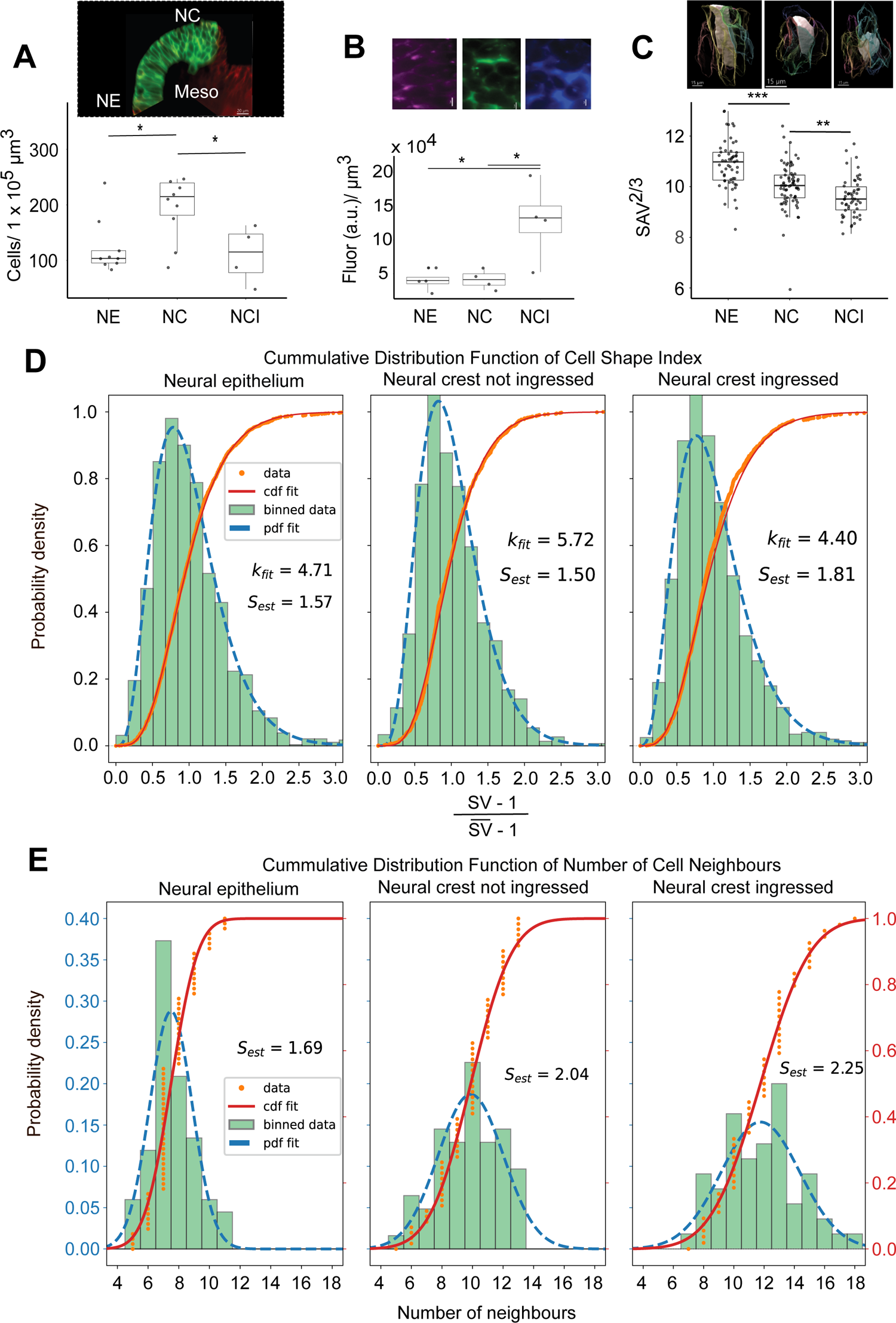
Geometric packing of neural crest cells during delamination. **A** Cell density of the neuroepithelium (NE), pre-ingressed neural crest (NC), and ingressed neural crest within mesoderm (Meso). Cells within the 3 regions of Wnt1Cre-^2 Tg/+^;mT/mG embryos were rendered using a semi-automated algorithm in IMARIS: n=8 embryos for NE, 10 embryos for NC, 4 embryos for NCI. ANOVA F(2,20)=5.63, p<0.05. Tukey HSD post hoc: NE–NC Tukey CD=-85.7, p<0.05; NC–NCI Tukey CD=99.0, p<0.05. **B** Fluorescence intensities of rhodamine-dextran dye in 1 × 10^6^ to 4 × 10^6^ µm^3^ optical tissue volumes within NE, NC and NCI were quantified in IMARIS n=3 embryos per region. ANOVA F(2,9)=7.83, p<0.05; Tukey HSD post-hoc: NE–NCI Tukey CD=-99061.0, p<0.05; NC–NCI Tukey CD=-86490.0, p<0.05. **C** Shape index of cells within the neural fold. Each dot represents a rendered cell. n=54 NE cells from 6 embryos, 73 NC cells from 5 embryos, and 55 ingressed NCI cells from 3 embryos. ANOVA F(2, 176)=30.9, p<0.05. Tukey HSD post-hoc: NE–NC Tukey CD=-0.83, p<0.001; NC–NCI Tukey CD=0.48, p<0.01. Insets represent clusters within each region highlighting a central cell (white) surrounded by its neighbours (in colour). **D** Distribution of the normalised cell shape indices represented by 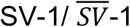 within the neural fold. 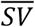 represents the mean shape index 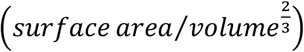 for each population. Estimated entropy (S_est_) was calculated by fitting the distribution of cell shape indices to a κ-gamma distribution. **E** Distribution of number of cell neighbours within the neural fold. Cells were segmented in 3D using IMARIS. Values for entropy (S_est_) were derived from fitting the number of neighbours to a normal distribution. n=67 NE cells from 4 embryos, 62 NC cells from 5 embryos, 55 NCI cells from 6 embryos. Wilcoxon test: NE– NC W=3344.5, p<0.001. NC–NCI W=957.5, p<0.001.

System entropies of collective granular materials^72^ and living cell clusters^71,73^ have been estimated based on the volumes and aspect ratios, respectively, of individual components that follow a k-gamma distribution. We adapted this statistical mechanics approach to estimate the entropies of the neuroepithelium, pre- and post-ingressed neural crest based on cell shape index and packing configurations. Using a nonlinear least squares method, we fit the cumulative distribution function to our data and found that cell shape indices closely follow a k-gamma distribution and numbers of cell neighbours follow a normal distribution in all 3 compartments. By both methods, estimated configurational entropy was highest among cells that had ingressed (Fig. 3D, E).

Neural crest cells exhibit cortical fluctuations (Mov. 10) that represent effective temperature in our model. To quantify these fluctuations, we used a transgenic FRET-based vinculin tension sensor (VinTS). This sensor links cortical actomyosin to adhesion receptors at the cell membrane and reports tension which is proportional to the average lifetime of the VinTS fluorophore^57^. VinTS was expressed conditionally in the neural crest using Wnt1:Cre2. Mean cortical tensions of pre-ingressed neural crest cells were significantly higher than those of neuroepithelial cells, and further elevated among ingressed cells (Fig. 4A, B). The amplitude of contractions, calculated as the deviation from the average lifetime across timepoints (root sum of squares, RSS), was also significantly higher among neural crest cells that had ingressed into mesoderm (Fig. 4C). By analogy to our simulation, these data suggest that effective temperature, like entropy, increases during cell ingression.

**Figure 4.**
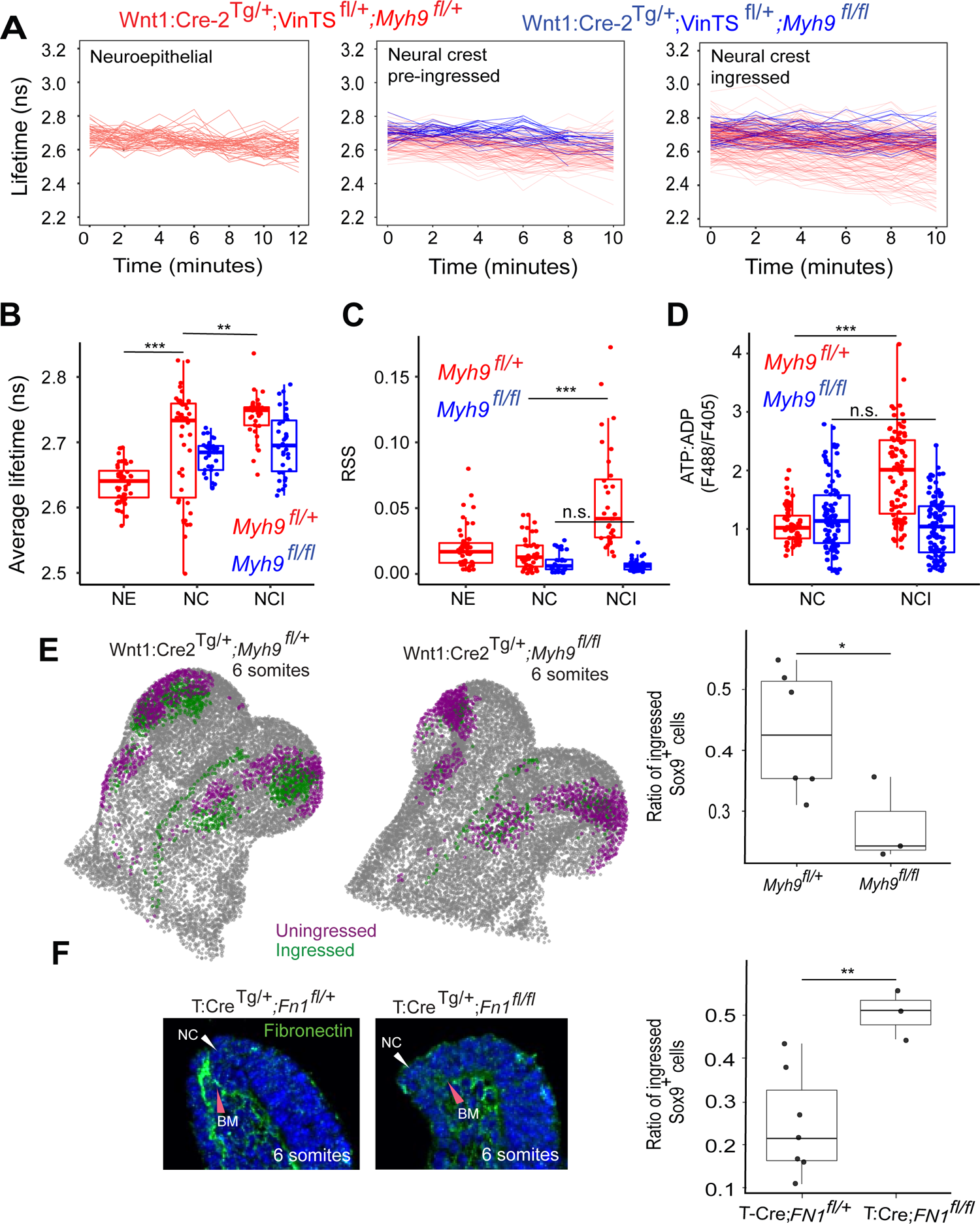
Manipulation of cortical fluctuation and basement membrane. **A** Cortical cell membrane fluctuations within the neural fold of Wnt1Cre-2^Tg/+^;VinTS^fl/+^;*Myh9*^*fl/+*^ (red curves) and Wnt1Cre-2^Tg/+^;VinTS^fl/+^;*Myh9*^*fl/fl*^ (blue curves) mouse embryos. Each trace is the lifetime fluorescence of VinTS over time and represents the cortical tension of individually segmented cells within the neuroepithelium (Wnt1Cre-2^Tg/+^; VinTS^fl/+^;*Myh9*^*fl/+*^ n=48 cells from 3 embryos), pre-ingressed neural crest (Wnt1Cre-2^Tg/+^; VinTS^fl/+^;*Myh9*^*fl/+*^ n=109 cells from 9 embryos; Wnt1Cre-2^Tg/+^;VinTS^fl/+^;*Myh9*^*fl/fl*^ n=34 cells from 3 embryos), and ingressed neural crest cells (Wnt1Cre-2^Tg/+^;VinTS^fl/+^;*Myh9*^*fl/+*^ n=182 cells from 11 embryos; Wnt1Cre-2^Tg/+^;VinTS^fl/+^;*Myh9*^*fl/fl*^ n=39 cells from 3 embryos). **B** Average cortical tension represented as VinTS lifetime values for cells in the neuroepithelium (NE), pre-ingressed neural crest (NC), and ingressed neural crest (NCI). Two-way ANOVA with significant main effect of cell position F(2, 183)=42.26, p<0.001; Tukey HSD post-hoc: NE–NC Tukey CD=-0.05, p<0.001, NC–NCI Tukey CD=0.03, p<0.01. The main effect of genotype was not significant F(1,183)=0.18, p=0.67. **C** Fluctuation amplitudes of each trace, calculated as the deviation (root sum of squares, RSS) from the average lifetime over time for cells within the NE, NC, and NCI. Two-way ANOVA with significant effect of interaction between cell position and genotype F(1,181)=41, p<0.001. Tukey HSD post-hoc: *Myh9*^*fl/+*^ NC–NCI Tukey CD=0.04, p<0.001; *Myh9*^*fl/fl*^ NC–NCI Tukey CD=-0.003, p>0.05. **D** ATP:ADP ratios of pre-ingressed, NC, and ingressed, NCI, neural crest cells of Wnt1Cre-2^Tg/+^;myrPercevalHR^fl/+^;*Myh9*^*fl/+*^ (red) and Wnt1Cre-2^Tg/+^;myrPercevalHR^fl/+^;*Myh9*^f*l/fl*^ (blue) mice. Each dot represents one cell. (Wnt1Cre-2^Tg/+^;myrPercevalHR^fl/+^;*Myh9*^*fl/+*^ n=56 NC cells from 4 embryos and n=87 NCI cells from 5 embryos; *Wnt1Cre-2*^*Tg/+*^*;myrPercevalHR*^*fl/+*^*;Myh9*^*fl/fl*^ n=98 NC cells from 7 embryos and n=106 NCI cells from 8 embryos). Two-way ANOVA with significant effect of interaction between cell position and genotype F(1,343)=69, p<0.001. Tukey HSD post-hoc: *Myh9*^*fl/+*^ NC–NCI Tukey CD=0.86, p<0.001; *Myh9* ^*fl/fl*^ NC–NCI Tukey CD=-0.20, p>0.05. **E** Proportion of ingressed cranial neural crest cells in intact, stage-matched 6-7 som Wnt1-Cre2^Tg/+^;*Myh9*^*fl/+*^ and Wnt1-Cre2^Tg/+^;*Myh9*^*fl/fl*^ embryos. Three-dimensional projections of nuclei identified using Nuclear Plotter. Purple label: Sox-9 positive nuclei of non-ingressed cells within the epithelium that are surrounded by high E-cadherin staining. Green label: Sox-9 positive nuclei deep to the epithelium surrounded by nil-low E-cadherin staining. Gray label: DAPI-stained nuclei. Right panel: Quantification of ingressed cells within intact embryos. Each dot represents one embryo: Wnt1-Cre2^Tg/+^;*Myh9*^*fl+l*^ n=6 embryos, *Myh9*^*fl/fl*^ n=3 embryos; t(6)=2.7, p<0.05. **F** Ingressed cranial neural crest cells in intact, stage-matched 4-7 som T:Cre;*FN*^*fl/+*^ and T:Cre;*FN*^*fl/fl*^ embryos. Whole-mount immunostaining for fibronectin in T:Cre;*FN*^*fl/+*^ and T:Cre;*FN*^*fl/fl*^ embryos. White arrowheads: neural crest (NC), red arrowheads: fibronectin-labelled basement membrane (BM). Right panel: Quantification of ingressed cranial neural crest cells in intact T:Cre;*FN*^*fl/+*^ and T:Cre;*FN*^*fl/fl*^. Each dot represents an embryo: T:Cre;*FN*^*fl/+*^ n=7, T:Cre;*FN*^*fl/fl*^ n=3; t(8)=-4.5, p<0.01.

To determine metabolic requirements associated with cortical fluctuations, we examined ATP:ADP ratios. It has been shown that glucose uptake is higher in migratory neural crest cells relative to pre-migratory cells in tissue explants^52^ and, *in vitro*, active cell migration correlates with high ATP:ADP ratios^74,75^. To assess this ratio *in vivo*, we generated a transgenic mouse strain, myrPercevalHR^fl/fl^, that harbours a floxed, myristoylated PercevalHR sensor^76^. *In vivo*, the sensor reported appropriately high or low ATP:ADP ratios in response to the presence or absence glucose and to glucose competition (Fig. S3A, B, Methods). Within the neural crest of Wnt1Cre2^Tg/+^;myrPercevalHR^fl/+^ embryos, ATP:ADP ratios were significantly higher among ingressed cells relative to pre-ingressed cells (Fig. 4D), indicating energy input is associated with the transition from one metastable state to another. We infer that energy alone is insufficient to allow the process to be reversible. Taking these observations together, we propose that raising effective temperature promotes the initial transition and increased entropy effectively traps cells within mesoderm.

### Manipulations of thermodynamic parameters *in vivo*

Our model predicts that decreasing cortical fluctuations would hamper the exit of cells from the epithelium. To test this hypothesis, we dampened cortical fluctuations by conditionally deleting *Myh9* that encodes the nonmuscle myosin IIA heavy chain, a regulator of the pulsatile nature of actomyosin contractions^77^. The mean tensions (Fig. 4A, B) and, especially, amplitudes of cortical fluctuations (Fig. 4C) were reduced in Wnt1Cre2^Tg/+^; VinTS^fl/+^;*Myh9*^*fl/fl*^ conditional mutants relative to *Myh9* heterozygous littermates. ATP:ADP ratios did not increase during ingression in mutant embryos (Fig. 4D), consistent with cortical fluctuations as an active process.

To determine the necessity of cortical fluctuations for cell ingression, we examined conditional *Myh9* mutants. Those mutants exhibited subtle gross dysmorphology (Fig. S4A) with an exaggerated neural crest bulge reflecting inappropriate pooling of cells (Fig. S4B). We measured the fraction of SOX9-positive neural crest cells that had ingressed in wholemount immuno-stained embryos ranging between 4 to 8 somite stages. Applying a custom script, we used E-cadherin levels^78^ and physical location to distinguish cells that had not ingressed within the epithelial neural crest from those that had ingressed within mesoderm (Methods). The proportion of ingressed cells was significantly reduced in *Myh9* mutant embryos (Fig. 4E), indicating that high amplitude cortical fluctuations are required for efficient cell ingression as predicted by our model.

The inability of neural crest cells to remodel the basement membrane has been shown to impair cell delamination and migration^31,32,79^. Those observations suggest the basement membrane represents a component of the system’s energy barrier. Based on our model, we expected that weakening the basement membrane would lower the energy barrier and enhance cell ingression. In mouse neural folds, immunostaining revealed that fibronectin (FN) is a significant component of the basement membrane and can be mostly eliminated by conditionally depleting its expression by mesodermal cells (TCre;*F1*^*fl/fl*^) (Fig. 4F). Quantification revealed that loss of fibronectin significantly increased the proportion of ingressed Sox9-positive cells in TCre;*Fn1*^*fl/fl*^ embryos, indicating that the height of the energy barrier negatively influences cell ingression as predicted by our model.

## DISCUSSION

Stochastic transitions between different states are central to multiple biological contexts^80-83^ such as cell fate changes^84-86^. Advances that equated active parameters to those of athermal systems have opened the possibility of examining thermodynamic properties of non-equilibrium, living systems^34,58,59,71-73^. By applying such an approach to test for a stochastic morphogenetic mechanism, this study complements others that have assessed thermodynamic parameters in whole organisms using calorimetry or actual temperature measurements^3,87-89^. The results here suggest that development is not merely dissipative^1-4,87-90^. Embryonic processes may have evolved to co-opt entropic tendencies that permit energy-efficient morphogenesis.

Cell ingression among vertebrates may have been adapted from ancient invertebrate cell behaviours observed at the archenteron margin in diploblasts^7,91,92^. As has been suggested, a small number of conserved physical rules may underlie basic morphogenetic motifs that were diversified by genetic variations^16,93^. Embryonic processes may have evolved through various combinations of energy-intensive order and energy conserving disorder. It may have been worthwhile to invest just enough energy to overcome an energy barrier to take advantage of transient disorder as long as it achieved a useful outcome. In the case of the neural crest, although various cues may regulate and vary the process in different organisms, the findings here suggest that cell ingression is a stochastic event at its core. Since a process like cell ingression is dissipative, the system does not revert to its original state, thereby stabilising new structures^94^.

## METHODS

### Animals

#### Mouse lines

Embryos from the following mouse lines were used in experiments: Wnt1Cre-2^Tg/+95^, Pax3Cre^96^, mT/mG, VinTS^fl/fl 57^, *Myh9fl/fl*^97^, and myrPercevalHR^fl/fl^ (detailed below).

#### Generation of myrPercevalHR^fl/fl^

PercevalHR is a fluorescent protein that changes its emission spectrum upon binding to ATP or ADP, thus providing a real-time measure of cell energy status^76^. To assess cellular metabolic flux *in vivo*, we generated a transgenic mouse line expressing a floxed, myristoylated^98^ PercevalHR sensor (*myrPercevalHR*^fl/fl^) that localises the reporter at the cell cortex. We then targeted the *Rosa26* locus in ES cells using vectors generated by Gateway cloning. Electroporation of vector constructs and generation of ES cell lines were conducted in a standard fashion by the Toronto Centre for Phenogenomics. Founders were outbred to CD1 mice to obtain an F1 generation through germline transmission.

Sensor function was validated *in vivo* by culturing T-Cre;*myrPercevalHR*^*fl/fl*^ embryos under conditions of glucose deprivation through glucose depletion or inhibition of glucose uptake (2-deoxyglucose treatment). ATP:ADP ratios were appropriately reduced in embryos that were incubated under these conditions (Supplementary Fig. 3).

### Imaging and analyses

#### Light-sheet microscopy and image processing

Three-dimensional images of the cranial region of embryonic day E8.5 embryos ranging from 4 to 8 somites were acquired using a Zeiss Lightsheet Z.1 microscope as previously described with modifications^57^. Embryos were embedded in a cylinder of 1% agarose in DMEM with 5% rat serum that was suspended in DMEM within the imaging chamber and kept at 37°C with 5% C0_2_. Time-lapse images were acquired every 3 minutes over a span of 2-3 hours per session. Raw, multiview images were aligned using Zen software and CZI files were imported into IMARIS (Oxford Instruments) for further analysis.

#### FLIM-FRET

Live embryos (E8.5) were imaged using the Nikon A1R laser scanning confocal microscope equipped with a PicoHarp 300 TCSPC module and a 400 nm pulsed diode laser (Picoquant) as described previously^57,99^. Live embryos were placed against the glass of the imaging chamber with 1% agarose gel composed of DMEM and 5% rat serum. Embryos were positioned with their sides against the glass to view the cranial region. The chamber was filled with DMEM and maintained at 37°C with 5% CO_2_. PTU files were read using FLIMfit^100^ and FLIMvivo^99^ (available at https://github.com/HopyanLab/FLIMvivo) to generate segmentation masks and fit the decay curve respectively.

#### Live confocal microscopy

To measure ATP:ADP ratios, live mouse embryos were embedded in agarose within imaging chambers maintained at 37°C with 5% CO_2_ as above and imaged using a Nikon A1R laser scanning confocal microscope. Samples were excited with 488 nm and 405 nm lasers for ATP and ADP, respectively, and emissions were captured using a 525/50 bandpass filter. The generated nd2 files were imported into Volocity (PerkinElmer) for analysis. For each file, the background signal was determined by defining 30 equal-sized regions of interest outside the tissue and within cell nuclei and subtracted from the ratios. The images were segmented manually to identify individual cells. Particles within each cell were weighted according to their areas. The ATP:ADP ratio for each cell was calculated as the sum of the weighted means of particles within the cell.

### 3D cell segmentation and neighbour counting

To examine three-dimensional cell shapes and neighbour relationships, CZI files were imported into IMARIS. Large files containing 3-dimensional, time lapse images were portioned into smaller sections to optimise computation. For each sub-section, 1 to 3 regions of interest were selected for cell segmentation. Cells were initially delineated using the membrane segmentation algorithm within IMARIS. Labelled cells were then inspected manually by examining them through *z* planes. Mislabelled cells, flagged as volumes less than 50 µm^3^ or greater than 1000 µm^3^, were appropriately fused and/or fragmented. Following this step, volumes that were still outside this range were discarded (Supplementary Fig. 2). Cell neighbours, defined as cells that shared an interface with the lead cell in 3D space, were counted manually.

### Quantification of ingressed vs. pre-ingressed cells

Fixed, whole embryos (E8.5, 4-8 somite stages) were immunostained against SOX9 (AB5535, mouse, EMD Millipore) and E-cadherin (610181, mouse, BD Biosciences), and imaged using a Nikon A1R laser scanning confocal microscope. The generated nd2 files were read and analysed using called Nuclear Plotter (available at https://github.com/HopyanLab/Nuclear_Plotter), a custom python script that identifies DAPI-labelled nuclei and allows users to classify cells based on nuclear and cellular immunostaining and proximity to designated tissue landmarks (e.g. neural crest). The location and counts of these nuclei are then exported into CSV files for analysis. The ingression state of SOX9-stained neural crest nuclei was distinguished according to intensity of E-cadherin signal and location. Pre-ingressed cells (shown as purple nuclei) were defined as cells within the epithelial layer that expressed high E-cadherin whereas cells within mesoderm with little to no E-cadherin were designated as ingressed (shown as green nuclei) (Fig. 4E). Ratios of ingressed cells were calculated as the fraction of ingressed cells divided by the total number of SOX9 positive nuclei.

### Vertex modelling and simulation

#### Two dimensional model of cell ingression

We generated a simple two-dimensional vertex model as a toy model to explore the effect of cortical fluctuations in the presence of distinct cell types that segregate. The model denotes energy as a function of cell areas and perimeters^101^

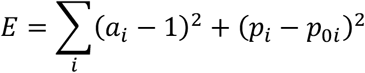

where the model parameter p_0i_ defines the target shape index for each cell and incorporates the relative strength of cortical tension and cell-cell adhesion. We added to this a segregation potential as a line tension between unlike cell types, providing an energy penalty for contacts between cells of different types to represent mismatched adhesion proteins.

Active cortical fluctuations were simulated by adjusting the shape index for each cell at each time step. To simulate the dissipative nature of the system, we applied gradient descent toward the local energy minimum at each step. The model parameters were treated as random variables drawn from cell type specific Gaussian distributions. As such, the model incorporates active and dissipative contributions to its dynamics.

The initial state involves two cell types that are segregated. We then modelled the onset of the conditions necessary for ingression by changing the line tensions of a few epithelial-type cells to be more like the mesodermal-type.

#### Effective temperature

Using the same model as above, we computed the “effective temperature” previously used in non-equilibrium granular systems^58,59^ as the ratio of particle diffusivity to susceptibility. If z defines the direction of cell motion (perpendicular to the interface between the two cell types represents epithelial cell ingression into mesoderm in our case), then the diffusivity is given by the mean square displacement over time.

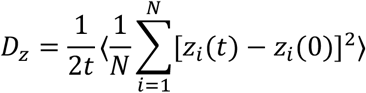

Susceptibility is the linear response to an external force (in the limit of small applied force).

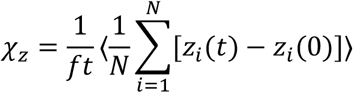

In the above formulae, N is the number of relevant cells, z_i_ is the position of the i^th^ cell, f is the applied force in the z-direction, and angled brackets denote the average over an ensemble of systems. We used an ensemble of 24 systems, each containing 132 cells with 12 migratory cells. Effective temperature is the ratio of diffusivity to susceptibility.

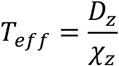

## CODE AVAILABILITY

https://github.com/HopyanLab/Ingression_Simulation

https://github.com/HopyanLab/Nuclear_Plotter

## AUTHOR CONTRIBUTIONS

C.P. co-developed the concept, performed embryonic imaging, quantitative analyses, generated *myrPercevalHR*^*fl/fl*^ transgenic mice, and wrote the manuscript; E.T. determined a scheme to assess thermodynamic parameters for the nonequilibrium neural crest, wrote custom scripts, performed all modelling and analysed some of the quantitative data; T.Y. performed quantitative assessments of cell ingression; M.Z. performed 3D magnetic tweezer measurement of tissue stiffness; H.T. provided technical support; Y.S. provided mechanical insights and funding; S.G. provided modelling insights and funding; S.H. co-developed the concept, assembled the collaborations, edited the manuscript, and provided funding.

## ACKNOWLEDGEMENTS

We thank Angie Griffin for animal husbandry, Kimberly Lau and Paul Paroutis for assistance in the Imaging Facility, Rudolph Winklbauer for discussions and review of the manuscript, Kelli Fenelon for discussions, R. Adelstein for permission to obtain Myh9^fl/fl^ mice from D. Schramek’s lab, and funding from the Canada First Research Excellence Fund (Medicine by Design Grand Questions Program, MbDGQ-2021-04) to S.G., Y.S. and S.H., and Canadian Institutes of Health Research (168992) to Y.S. and S.H..

## SUPPLEMENTARY MOVIES

**Supplementary movie 1**

Neural crest cells (green) are seen at the dorsal lips of the closing neural fold and migrating ventrally through mesoderm (red) of a Wnt:Cre2^Tg/+^; mT/mG embryo at 12 somite stage.

**Supplementary movie 2**

Cells ingress at different points along the cephalic neural crest. Representative cell surface-rendered cranial neural crest within the midbrain of a Wnt:Cre2^Tg/+^;mT/mG embryo at 4 somite stage showing the initial group of ingressing neural crest cells (solid colours) and their adjacent pre-ingressed neighbours (green).

**Supplementary movie 3**

Apical view of a neural crest cell ingressing at 4 somite stage. The apical region of the central cell (yellow) diminishes relative to its immediate neighbours as it leaves the plane, similar to a T2 transition.

**Supplementary movie 4**

Lateral view of neural crest cell ingression. Cell surface rendering demonstrates an ingressing cell (solid white) undergoing shape fluctuations as it moves to a more basal position relative to its immediate neighbours (green) within a Wnt:Cre2^Tg/+^;mT/mG embryo (5 somites).

**Supplementary movie 5**

Lateral view of a surface-rendered chain, or ‘stalactite’ of ingressing neural crest cells comprised of a lead cell (yellow) tightly associated with follower cells within the midbrain of a Wnt:Cre2^Tg/+^;mT/mG embryo (5 somites). Marked cell shape fluctuations are observed.

**Supplementary movie 6**

Individual streams, or ‘stalactites’, of neural crest cells coalescing with the mesoderm of a Wnt:Cre2^Tg/+^;mT/mG embryo (7 somites).

**Supplementary movie 7**

Ingressed and coalesced neural crest cells (green) within mesoderm (red) at the midbrain are observed remodelling their group into a narrower configuration prior to deeper migration in a Wnt:Cre2^Tg/+^;mT/mG embryo (8 somites).

**Supplementary movie 8**

Vertex model simulation of neural crest cells (green) within neuroepithelium (blue) and the underlying mesoderm (red). When simulated with low amplitude (6%) shape fluctuations, neural crest cells spontaneously move incompletely toward mesoderm with some remaining within the epithelial layer despite having a high affinity (60% - represented as line tension) for mesoderm.

**Supplementary movie 9**

Vertex model simulation of neural crest cells (green) within neuroepithelium (blue) and the underlying mesoderm (red). When simulated with high amplitude (16%) shape fluctuations, neural crest cells move completely into mesoderm with some remaining within the epithelial layer.

**Supplementary movie 10**

Cell surface rendering of an ingressing neural crest cell within the midbrain of a Wnt:Cre2^Tg/+^;mT/mG embryo (5 somites) demonstrates marked shape fluctuations.

**Extended Data Figure 1.**
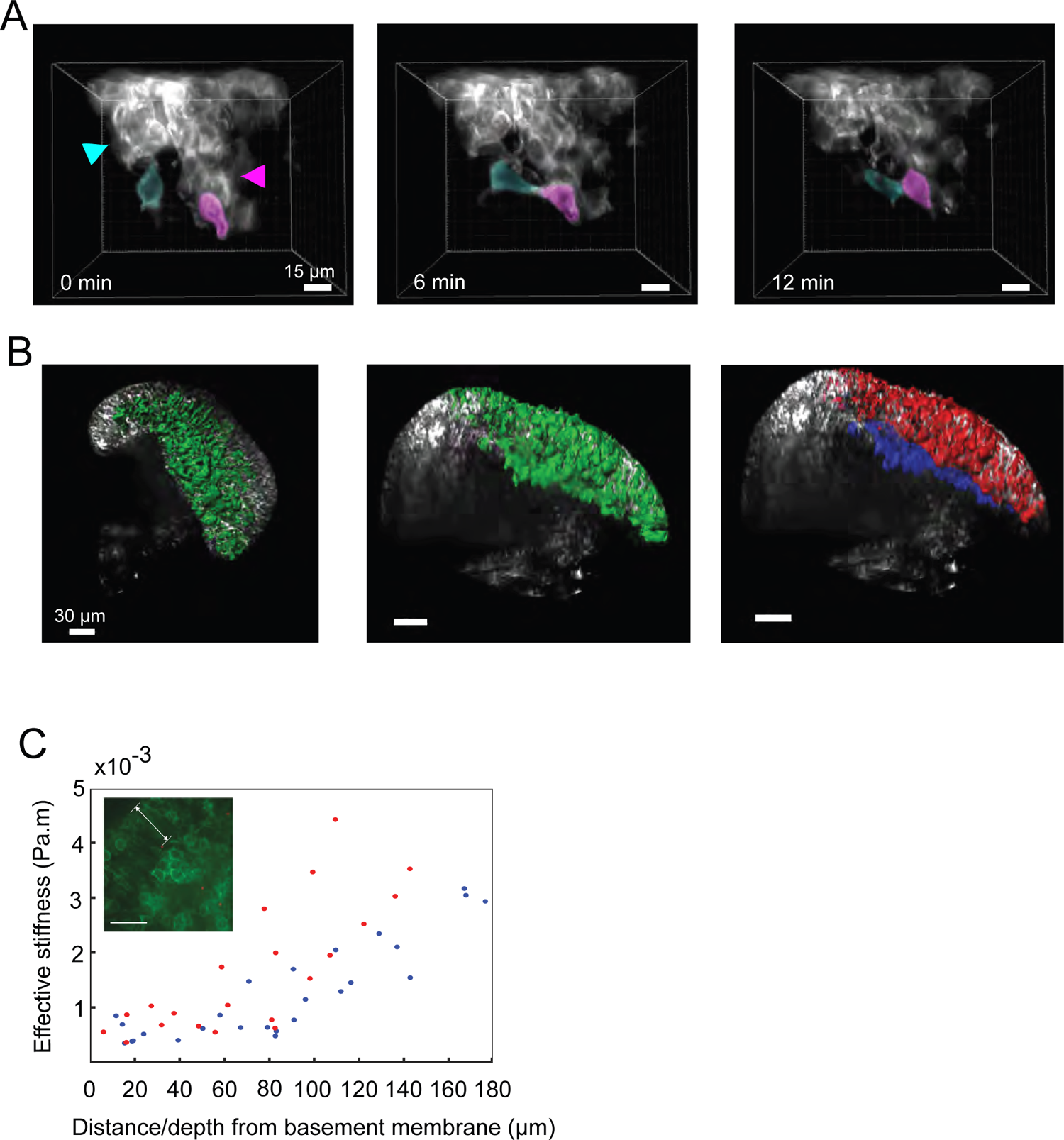
**A** Still images from Supplementary Movie 2 showing lead cells (blue and pink) of adjacent ingressing “stalactites” (blue and pink arrowheads) coalescing after ingression. Scale bars represent 15 µm. **B** Neural crest cells (green) in the midbrain of a 7 somite Pax3:Cre;mT/mG embryo showing pre-ingressed cells in the epithelium (red) and coalesced group of ingressed cells (blue). Scale bars represent 50 µm. **C** Tissue stiffness within the neural fold. Stiffness was measured by injecting magnetic beads (red dots in inset) into the neural crest and underlying mesoderm of mouse embryos (n=3 embryos ranging from 6-8 somite stage) and measuring their displacement within a uniform magnetic field gradient. White double arrow line in inset signifies the epithelium.

**Extended Data Figure 2.**
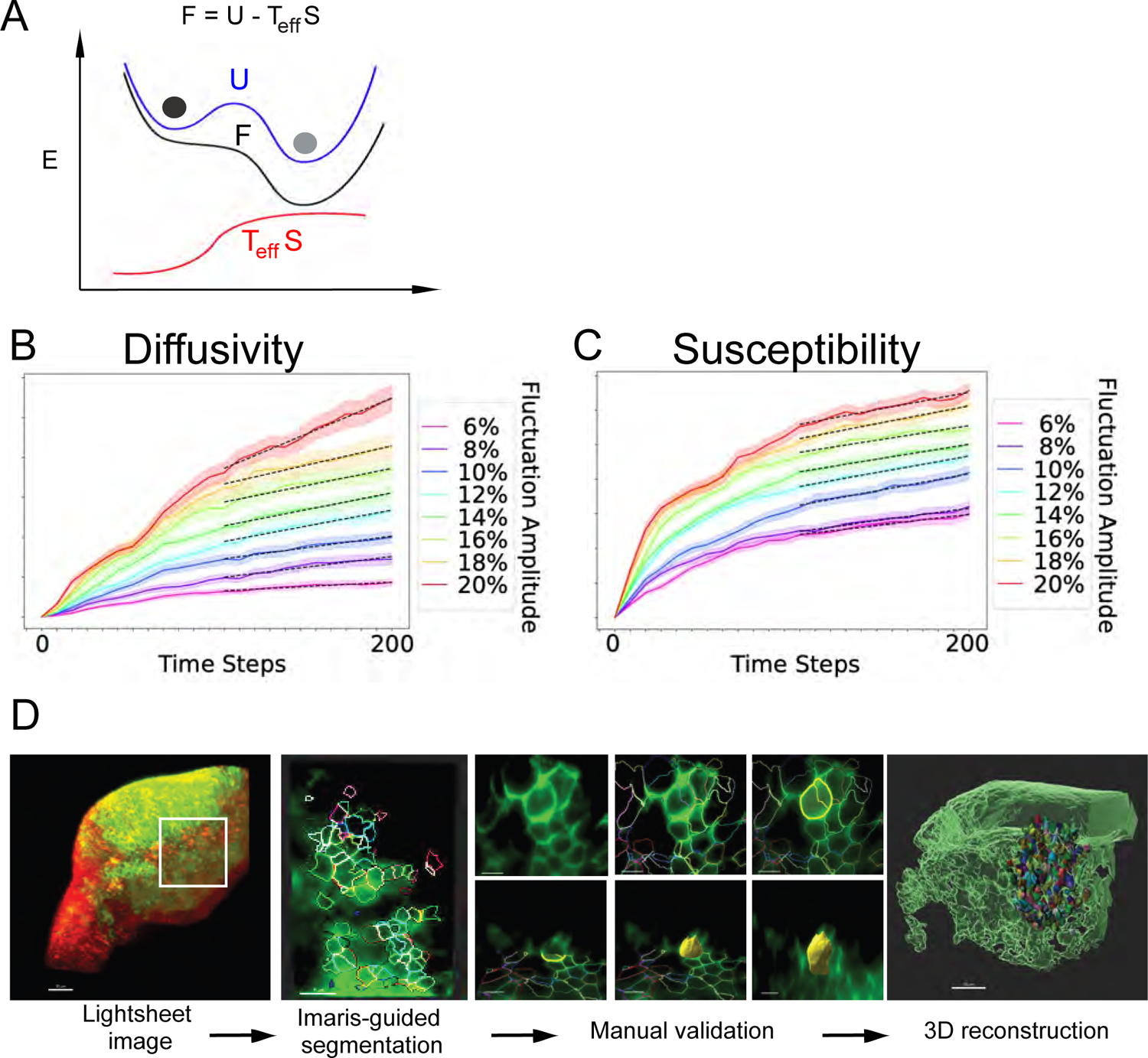
**A** Hypothetical energy landscape (blue curve) a neural crest cell (black circle) would follow to reach another metastable state within mesoderm (grey circle). In principle according to the Helmholtz free energy (F) equation, a cell can overcome an internal energy barrier (ΔU) if the entropic term (effective temperature T_eff_ * entropy S) is high. An increase in the entropic term (red curve) would diminish free energy, resulting in a directional preference (black curve) that renders cell ingression thermodynamically favourable. **B** In the vertex model, cell diffusion as a function of time becomes linear at later time steps. We fit the linear portion of the curve to calculate slope with respect to time. **C** Susceptibility to a small force (10^−4^) also becomes linear at later time steps, and the slope of the linear portion was calculated. Effective temperature was calculated by taking the ratio of the two slopes as per the Fluctuation-Dissipation Relation D ∼ T_eff_ χ. For A and B, solid colour curves represent means and shading represents standard error (sigma/square root(n)) of 48 simulations. **D** Workflow for semi-automated cell segmentation. i Using IMARIS, lightsheet images of intact Wnt1:Cre2^Tg/+^;mT/mG embryos were partitioned into smaller volumes for further processing and analysis. ii Cell boundaries were identified using fluorescence thresholds in IMARIS. iii Segmented cell membranes were validated by manual inspection of image slices in xy and yz planes. Relatively small (< 200 µm^3^) and large (> 1000 µm^3^) segments were re-evaluated and fused or fragmented as appropriate. Segmented volumes at the extremes that were less than 50 µm (unrealistic for a cell) or greater than 2000 µm^3^ (multiple cells) were deleted. iv Validated cells were reconstructed in 3D for analysis of cell behaviour and neighbour relationships.

**Extended Data Figure 3.**
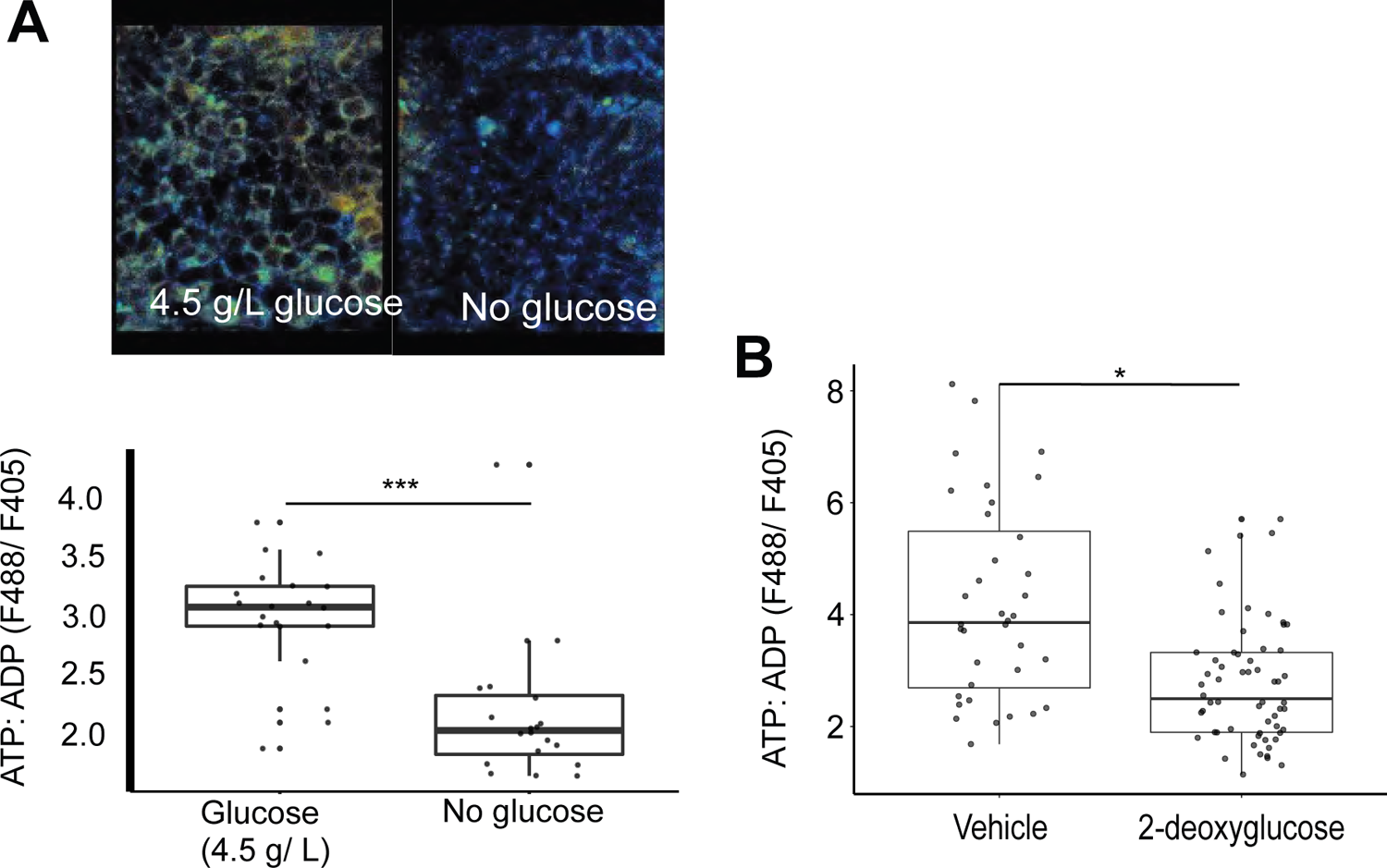
Validation of transgenic myrPercevalHR sensor *in vivo*. **A** E8.5 T:Cre;myrPercevalHR^fl/+^ embryos were cultured in DMEM + 2% rat serum, with or without 4.5 g/L glucose for 30 minutes. ATP:ADP ratios were calculated and corrected for background fluorescence following excitation at 488 nm or 405 nm. n=20 cells from 2 embryos for each condition; t-test p=3.258 ×10^−5^. **B** T:Cre;myrPercevalHR ^fl/+^embryos (E8.5) were cultured in regular medium (DMEM) with vehicle or 2-deoxyglucose (2 mM) for 30 minutes. n=60 cells from 3 embryos cultured in vehicle, 100 cells from 3 embryos cultured in 2-deoxyglucose; p=0.029, t-test.

**Extended Data Figure 4.**
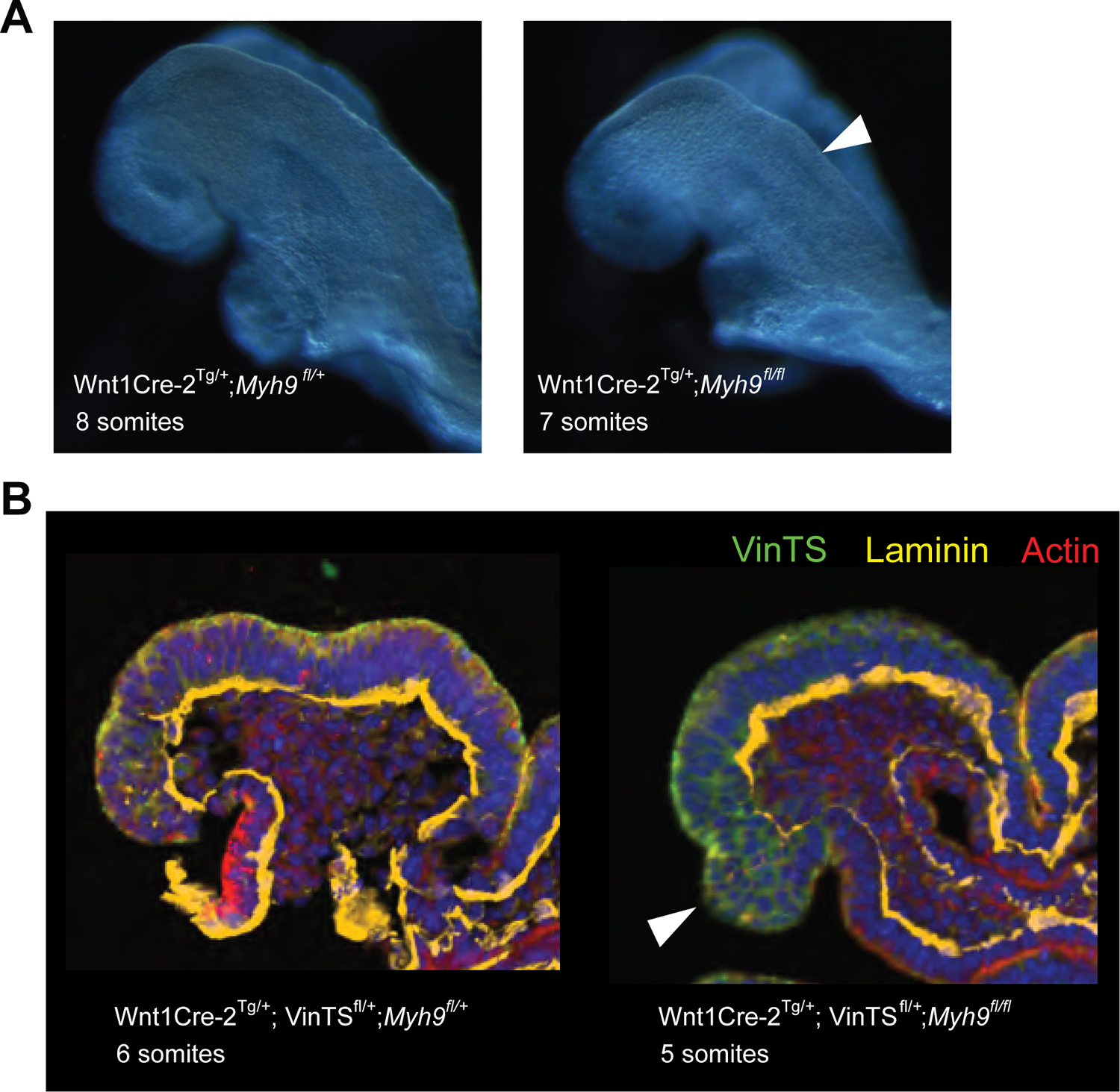
Conditional knockout of Myh9 induces a mild phenotype at the neural crest. **A** Relative to littermate controls, the neural folds of mutant Wnt1:Cre-2 ^Tg/+^;*Myh9* ^*fl/fl*^ embryos exhibited a subtly altered curvature (white arrowhead). **B** Immuno-stained midbrain sections of Wnt1Cre-2 ^Tg/+^;*Myh9* ^*fl/+*^(wild type) and Wnt1Cre-2 ^Tg/+^; *Myh9* ^*fl/fl*^ (conditional mutant) embryos. White arrowhead points to an inappropriate accumulation of cells at the neural crest.

